# Eye-hand coordination during online reach corrections is task-dependent

**DOI:** 10.1101/2021.06.13.448238

**Authors:** Anouk J. de Brouwer, Miriam Spering

**Affiliations:** Department of Ophthalmology & Visual Sciences, University of British Columbia, Vancouver, BC, Canada; Institute for Computing, Information and Cognitive Systems, University of British Columbia, Vancouver, BC, Canada; Djavad Mowafaghian Centre for Brain Health, University of British Columbia, Vancouver, BC, Canada

**Keywords:** online control, visuomotor, perturbation, reaction time, saccade

## Abstract

To produce accurate movements, the human motor system needs to deal with errors that can occur due to inherent noise, changes in the body, or disturbances in the environment. Here, we investigated the temporal coordination of rapid corrections of the eye and hand in response to a change in visual target location during the movement. In addition to a ‘classic’ double-step task in which the target stepped to a new position, participants performed a set of modified double-step tasks in which the change in movement goal was indicated by the appearance of an additional target, or by a spatial or symbolic cue. We found that both the absolute correction latencies of the eye and hand and the relative eye-hand correction latencies were dependent on the visual characteristics of the target change, with increasingly longer latencies in tasks that required more visual and cognitive processing. Typically, the hand started correcting slightly earlier than the eye, especially when the target change was indicated by a symbolic cue, and in conditions where visual feedback of the hand position was provided during the reach. Our results indicate that the oculomotor and limb-motor system can be differentially influenced by processing requirements of the task and emphasize that temporal eye-hand coordination is flexible rather than rigid.

**New & Noteworthy:** Eye movements support hand movements in many situations. Here we used variations of a double-step task to investigate temporal coupling of corrective hand and eye movements in response to target displacements. Correction latency coupling depended on the visual and cognitive processing demands of the task. The hand started correcting before the eye, especially when the task required decoding a symbolic cue. These findings highlight the flexibility and task-dependency of eye-hand coordination.

## Introduction

How humans adjust and optimize movements to correct for errors that are due to sensory and motor noise, changes in the body, or external disturbances is a major focus of current neuroscience research. In the laboratory, tasks that artificially produce movement errors have revealed the ability to rapidly correct for errors during the movement (i.e., movement corrections), as well as the ability to adjust the movement to consistent errors over the course of several repetitions (i.e., motor adaptation). Here, we investigate the temporal coordination of movement corrections in eye and hand movements in a reaching task.

The significant progress in understanding how the sensorimotor system corrects for errors in eye and hand movements is largely based on studies that have investigated these two motor systems separately. The double-step task has been widely used to study both movement corrections and motor adaptation in eye and hand movements. In this classic paradigm, the visual target is displaced at the time of movement onset to simulate a spatial error. In the case of saccadic eye movements, the endpoint error after the saccade toward the initial target location triggers a second, corrective saccade to the new target location (Becker and Jürgens 1979; Hallett and Lightstone 1976; Joiner et al. 2010; Tian et al. 2013). If the target is repeatedly displaced to the same location, motor adaptation will result in an adjustment of the initial saccade (Herman et al. 2013; McLaughlin 1967; Tian et al. 2009). The double-step paradigm has also been used extensively to investigate corrections of reach movements, and a few studies have used this paradigm to investigate adaptation of reach movements (e.g., Magescas and Prablanc 2006). Because the duration of reach movements is much longer than that of saccades, movement corrections in response to a displacement of the reach target typically occur online (i.e., ‘in flight’) (Georgopoulos et al. 1981; Goodale et al. 1986; Megaw 1974; Soechting and Lacquaniti 1983) with reaction times as short as ~110 ms (Brenner and Smeets 1997; Day and Lyon 2000; Prablanc and Martin 1992). Another popular paradigm to study reach corrections is the cursor displacement paradigm, in which hand movements correct for a perceived deviation of the reach trajectory with short latencies (Brenner and Smeets 2003; Franklin and Wolpert 2008; Sarlegna et al. 2003; Saunders and Knill 2003). Together, these findings indicate that visual information is continuously used to control and correct movements.

The few studies that simultaneously measured both eye and hand movements in response to a displacement of a visual target have made several interesting observations. First, hand movement corrections start earlier when a corrective saccade accompanies the reach, as opposed to when the eyes are instructed to fixate (Abekawa et al. 2014; Diedrichsen et al. 2004), showing a facilitative effect of eye movements on hand movements. Second, whereas the eye usually leads the hand by (several) hundred milliseconds when initiating a goal-directed reach (e.g., Land and Hayhoe 2001; Prablanc et al. 1979), the hand might start correcting for spatial errors before the eye (Abekawa et al. 2014; Gritsenko et al. 2009), although the opposite finding (i.e., the eye corrects before the hand) has also been reported (Neggers and Bekkering 2002).

The tasks used to investigate movement corrections of the eye and/or hand are often limited to simple target step paradigms. For hand movements, it has been shown that the visual characteristics of the target (Kozak et al. 2019; Veerman et al. 2008), as well as the presence of visual distractors (Reichenbach et al. 2014) can influence the duration within which corrections are initiated, but eye movements were either not measured (Veerman et al. 2008) or participants were instructed to fixate (Reichenbach et al. 2014) in these studies. To thoroughly investigate the temporal coordination of eye and hand movement corrections, we assessed the timing of corrections in various stimulus and task conditions.

On one hand, we might expect a tight temporal coupling of corrective eye and hand movements, independent of stimulus and task conditions, as a result of shared visual processing and computation of the required correction for the eye and the hand. There is ample evidence for a close behavioral and neurophysiological connection of both movements (de Brouwer et al. 2021; Gopal and Murthy 2015; Land and Hayhoe 2001; Prablanc et al. 1979). On the other hand, temporal coupling might be loose, or perhaps variable across conditions, in order to optimize performance. Loose temporal coupling could result from independent visual processing and/or independent initiation of the required correction. For example, several studies have revealed active inhibition of saccades during the execution of hand movements (Mrotek and Soechting 2007; Neggers and Bekkering 2000), presumably to prevent a temporary distortion of the retinal image by eye movements (Ross et al. 2001). When correcting for a target displacement, it might be optimal to maintain fixation at the original target and perform a reach correction in peripheral vision before making a corrective saccade. One could also argue the opposite, namely, that it is beneficial to make a saccade to the new target as soon as possible. Recent work has shown that errors in the reach trajectory (i.e., due to a cursor jump) evoke the earliest and most vigorous corrections when gaze is directed at the reach target, as compared to surrounding locations (de Brouwer et al. 2018)(see also Abekawa et al. 2014; Diedrichsen et al. 2004). As such, making a saccade to the new target as early as possible might be particularly beneficial when peripheral visual feedback of the hand can be used to improve reach accuracy (i.e., when the hand is visible) as opposed to when this feedback is not available. This could result in differences in temporal coupling in conditions with and without feedback.

Participants performed a set of tasks in which they were asked to reach, as rapidly and accurately as possible, toward a visual target using a robotic manipulandum, while their hand and eye movements were recorded. In a subset of trials, an unpredictable change in the movement goal occurred. We included a classic double-step task (Hallett and Lightstone 1976), in which the target ‘stepped’ to a new location, triggering an immediate, exogenously driven saccade to that location. In addition, we designed a set of modified double-step tasks with the intention to manipulate the latency of the saccade to the new movement goal. In these tasks, the change in movement goal was indicated by the appearance of an additional target, or by a spatial or symbolic cue, placing different demands on visual and cognitive processing. Whereas the visual target displacement could be expected to trigger an immediate saccade, the additional target and cue conditions were designed to prevent triggering an immediate saccade and instead produce a later, more voluntary (or endogenously driven) saccade. Thus, we predicted that our modified double-step tasks would delay the corrective saccade to the new goal location. We tested the hypothesis that reach corrections would be delayed to the same extent as corrective saccades (strong temporal coordination), resulting in a constant relative latency between eye and hand corrections. In addition to varying the visual characteristics of the target change, we manipulated the presence of visual feedback of the hand. We hypothesized that the presence of hand feedback would speed up the corrective saccade to allow optimal monitoring of the reach trajectory (de Brouwer et al. 2018), and tested whether this also sped up the reach correction (Reichenbach et al. 2009). Overall, we aim to contribute to a better understanding of the temporal coordination of eye and hand movements during online reach corrections, the role of visual and cognitive demands in different task contexts, and the role of visual feedback.

## Methods

### Participants

Nineteen participants (mean age 26 years, range 19-36 years, 5 female) completed the experiment and were compensated $12/hour for their participation. Three other participants did not complete the experiment because the experimenter could not achieve sufficiently accurate calibration of the eye tracker (mean error <1.5° and maximum error <3.0° visual angle). All participants were self-reported right-handed and had normal or corrected-to-normal visual acuity. The study was approved by the University of British Columbia Behavioural Research Ethics Board. Participants provided written informed consent before the start of the experiment.

Of the participants who completed the experiment, three participants were excluded from the analysis because they had less than 4 (out of 10) valid trials for any combination of task × target change time × target change direction in two or more tasks. The most common reason for an insufficient number of valid trials was that an initial saccade to the target could not be detected, likely due to eye tracking difficulties. This exclusion resulted in 16 complete and analyzed data sets for the experiment.

### Experimental setup

Participants performed reaching movements to visual targets with their right hand. They were seated in a chair with their forehead resting against a pad and their hand holding on to the handle of a robotic manipulandum that moves in the horizontal plane (KINARM End-Point Robot; BKIN Technologies Ltd., Kingston, ON, Canada). A mirror was mounted horizontally above the handle, and an LCD monitor was mounted horizontally above the mirror, such that stimuli were projected in the plane of the handle when looking onto the mirror while the view of the arm was blocked (Fig. 1A). Kinematics of the handle were recorded with a temporal resolution of 1129 Hz, resampled to 1000 Hz. Eye movements of the left eye were recorded using a built-in monocular video-based eye tracker (Eyelink 1000, SR Research Ltd., Kanata, ON, Canada) with a temporal resolution of 500 Hz, resampled to 1000 Hz. The pupil was detected with a proprietary algorithm that accounted for small head movements, which were measured using a target sticker placed on the participant’s forehead or cheek. The eye tracker was calibrated for the 2D horizontal workspace using proprietary algorithms (BKIN Technologies).

**Figure 1.**
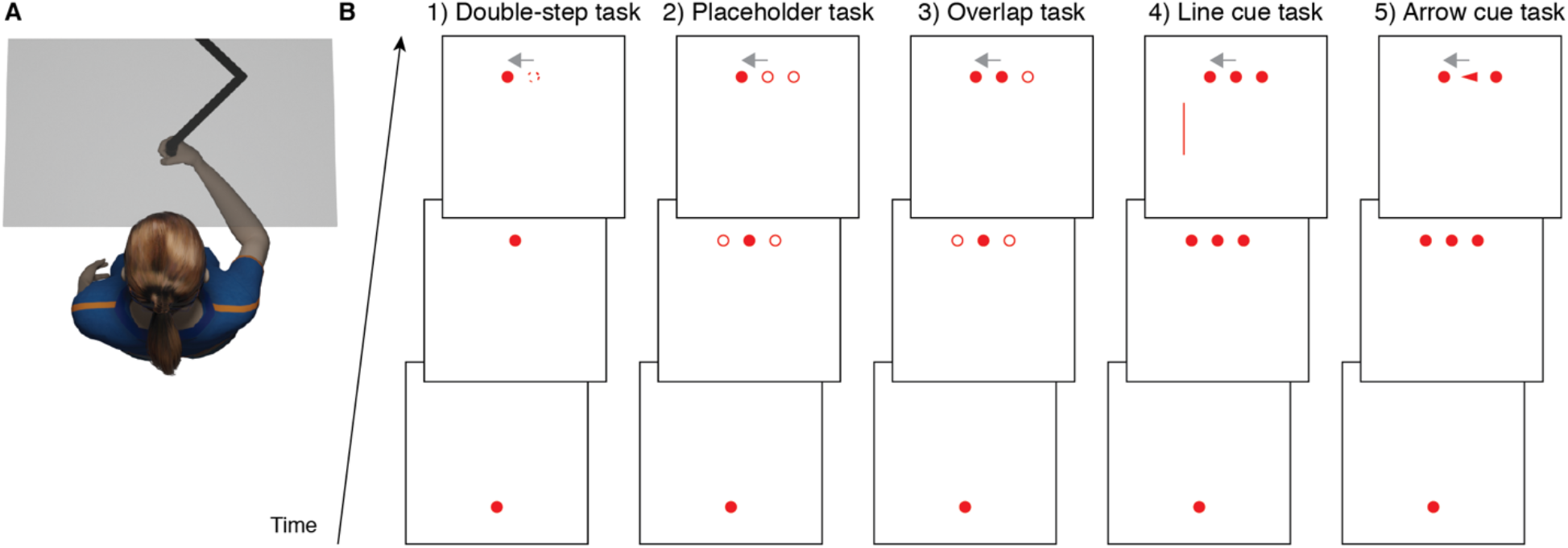
Experimental setup and tasks. A) Participants performed reaching movements in the horizontal plane using a KINARM robotic manipulandum. Vision of the hand was blocked by a mirror onto which the stimuli were projected such that they appeared in the plane of the handle. The mirror is depicted as transparent to illustrate the location of the hand and the manipulandum. B) In each task, participants moved their hand from a start position to one of three target positions 20 cm in front of the start position (both shown as red circles). In a subset of trials, the movement goal changed from the central position to a position 5 cm to the left or right of the central position (shown by the grey arrow pointing to the left that was not visible to the participant). In the double-step task (1), the target stepped from the central to the left/right position (the dotted circle indicates the original, invisible target position). The placeholder task (2) was identical to the double-step task, except that the two alternative target positions were indicated with open circles together with the target position. The overlap task (3) was identical to the placeholder task, except that the original target position remained ‘filled in’ when the target jumped to its new position. In the line cue task (4), all three possible target locations were indicated, and the (new) target was indicated by a spatial cue: a line on the left for the left target, a line on the right for the right target, no line for the central target). In the arrow cue task (5), the (new) target was indicated by a symbolic cue location at the central target: a leftward pointing arrow for the left target, a rightward pointing arrow for the right target, a circle for the central target.

### Task and visual stimuli

Figure 1B provides a schematic illustration of the tasks. The hand position was represented on the screen as a cursor (1 cm diameter white circle) aligned with the handle. All other stimuli were presented in red (5.9 cd/m^2^) on a black background (0.9 cd/m^2^) for clear, high-contrast visibility. Each trial began with the presentation of a start position (2 cm diameter circle) at the horizontal center of the screen near the participant. Participants were instructed to move the cursor and their gaze to the start position to initiate the trial. The cursor had to be at the start position for 250 ms, and the recorded eye position on the screen had to be within a 5 cm radius of the start position at the end of the 250-ms period. After a random delay of 250 to 500 ms, the start position disappeared and the reach target (2 cm diameter circle) appeared 20 cm in front of the start position and in line with the start position or 5 cm to the left or right of the start position. Participants were instructed to reach toward the target as quickly and accurately as possible. In 40% of trials, the movement goal remained unchanged during the trial. In the remaining 60% of trials, a sudden change in movement goal from the central to either the left or right target location was presented. Targets that initially appeared on the left or the right never changed location, but were included to discourage participants from initiating anticipatory movements to the central target. If the target changed, participants were instructed to move their hand to the new target as soon as possible. The target disappeared 250 ms after detection of the reach offset, defined by the velocity of the handle falling below a threshold of 2 cm/s for 250 ms. To encourage participants to perform fast reaching movements throughout the experiment, a message ‘Too slow’ was displayed on the screen if the movement time was longer than 800 ms in unperturbed trials, or longer than 1100 ms in trials with a change in movement goal. A new trial started after a 500 ms inter-trial-interval.

Participants performed five versions of the main task, in separate blocks (Fig. 1B). (1) In the classic double-step task, a single reach target stepped from the central to the left or right target position. (2) In the placeholder task, placeholders (2 cm diameter open circles) were presented in addition to the target at the two non-target locations. When the target stepped, it would ‘swap’ position with one of the placeholders. (3) The overlap task was similar to the placeholder task, except that when the target stepped, the central circle would remain filled (i.e., the old target would stay in its position when the new target was presented). (4,5) In the line and arrow cue tasks, three filled targets were presented at the three target locations. (4) In the line (i.e., spatial) cue task, a reach to the left or right target, or a change in movement goal from the central to the left or right target, was indicated by the appearance of a vertical line (10 cm × 0.2 cm) presented 10 cm to the left or right of the midline, and vertically centered between the start and target position. (5) In the arrow (i.e., symbolic) cue task, a reach or change to the left or right target was indicated by the central circle changing into a leftward or rightward pointing triangle (2.5 × 1.5 cm).

To ensure that the time of the change was unpredictable and that our results were not determined by the time of the target change, the target jump or appearance of the cue was triggered by one of three events: the onset of the saccade (*y* gaze position reached a third of the distance between the start and target position), the onset of the reach movement (the cursor had fully moved out of the start position), or the time where the handle passed the midpoint between the start position and the target position. The execution of the saccade and the onset of the reach movement are commonly used target change triggers in target displacement tasks (Diedrichsen et al. 2004; Gritsenko et al. 2009; Oostwoud Wijdenes et al. 2013; e.g., Prablanc and Martin 1992). We included the late target change time to make sure that the timing of the corrective saccade would not be influenced by the saccade refractory period, which is commonly observed to be around 150 ms (Carpenter 1977).

### Procedure

Each participant performed one practice block and ten experimental blocks, with short breaks in between blocks. Half of the experimental blocks were performed with online visual feedback of the hand position during the reach, and the other half was performed with endpoint feedback only. In the latter case, the cursor disappeared when the reach target was presented and reappeared after movement offset. The order of tasks and feedback was counterbalanced. Twenty different orders for the five tasks were created by starting with a 5×5 Latin Square in which every task occurs once in each row and once in each column, and then duplicating this square three times while randomly permuting the columns. Half of the participants performed every task first with online cursor feedback and then with endpoint cursor feedback, while the other half performed every task first with endpoint cursor feedback and then with online cursor feedback. The eye tracker was typically recalibrated at the start of each block, however, in some cases where calibration proved difficult, the eye tracker was recalibrated every two blocks. No instructions on eye movements were given except for directing the eyes to the start position at the beginning of the trial. The practice block consisted of 20 trials without a target change. Each experimental block consisted of 20 repetitions of unperturbed trials to the central target, 10 repetitions of unperturbed trials to the left/right target, 10 repetitions of trials in which the movement goal changed to the left/right during the saccade to the target, 10 repetitions of trials in which the movement goal changed to the left/right target following reach onset, and 10 repetitions of trials in which the movement goal changed to the left/right following the handle passing the point midway between the start and target position, resulting in a total of 100 trials per block. Participants took about 90 minutes to complete the experiment.

### Data analysis

The timing of the appearance of all visual stimuli and events (start position, target, target displacement, cue) was corrected offline by the delay in the system (57 ms), which was measured using a photodiode after the completion of data collection. Trials were excluded if the offline analysis showed that the target change had occurred before the event that triggered it (<0.5% of trials).

#### Hand and eye movement data preprocessing

The *x* and *y* positions of the center of the handle in the horizontal plane were used for the analysis of hand movements. For each trial, the onset of the reach was defined as the first moment in time where the resultant velocity of the handle was greater than 5 cm/s. In the offline analysis, the reach offset was defined as the first moment in time where the resultant velocity of the handle was smaller than 5 cm/s for 250 ms after the handle had passed the point midway between the start and target position. Trials were discarded if the onset of the reach occurred before the target appeared (1% of trials), or more than 1000 ms after the target appeared (<0.5%). Trials were also discarded if the *y* amplitude of the reach was less than half the target distance, or the *x* amplitude was less than half of the target displacement in change trials (1%).

The *x* and *y* eye positions on the screen, as obtained from the calibrated eye tracker, were used for analysis of the eye movements. Eye blinks, intervals in which the pupil signal was missing, and intervals in which the eye was detected outside the screen were removed from the *x* and *y* eye positions. For intervals with missing data for less than 100 ms (i.e., shorter than a typical blink duration), *x* and *y* eye position were linearly interpolated. Eye positions were filtered with a 2nd order lowpass Butterworth filter with a cut-off frequency of 30 Hz. *X* and *y* eye velocity 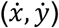 were computed by numerically differentiating *x* and *y* eye position signals.

Next, 2D eye positions on the screen were converted to an eye-based 3D coordinate system (Singh, Perry & Herter, 2016), assuming that the height of the stimulus plane relative to the eye is fixed (i.e., *z* is constant). The 3D eye positions on the screen were then transformed to an eye-based spherical coordinate system according to

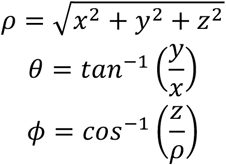

with *ρ* the radial distance from the eye to the point of gaze on the screen, *θ* the azimuthal angle in the *xy*-plane, and *ϕ* the elevation angle. Next, *ρ*, *θ*, and *ϕ* were differentiated to obtain 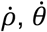 and 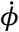:

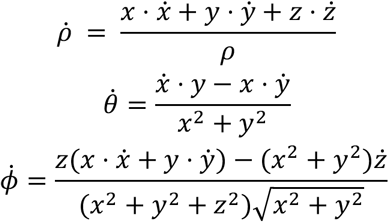

Note that 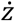 is assumed to be zero, reducing some parts of the equations to zero. Finally, the eye angular velocity *ω* was calculated according to

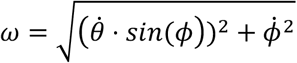

Saccades were detected by searching for intervals in which angular eye velocity was larger than 100°/s for at least 4 ms. Next, saccade onset was defined as the last sample in which eye velocity was below a threshold of 30°/s for 2 ms. Saccade offset was defined as the first sample in which eye velocity fell below a threshold of 30°/s for 2 ms.

Target saccades were defined as saccades that started between 50 ms before (predictive saccades) and 1000 ms after target appearance, with a *y* onset position before the midpoint between start and target position, and a minimum *y* amplitude of half the target distance. Trials were discarded if a target saccade was not detected (4% of trials). Corrective saccades from the initial to the final target were defined as saccades that started at least 50 ms after the target change, with a *y* onset and offset position past the midpoint between start and target position, and a minimum *x* amplitude of half the target displacement. Change trials were discarded if a corrective saccade was not detected (1% of trials). 93% of trials were included in the analysis.

#### Hand and eye movement analysis

We obtained the reaction times of the initial saccade and reach movements to the target as described above, using a velocity threshold of 30°/s to define the onset of the saccade, and a threshold of 5 cm/s to define the onset of the reach. We then computed, for target change trials, the latencies of the corrective responses of the eye and hand. The latency of the corrective saccade was defined as the latency of the first saccade directed towards the new target position following the target change. The latency of the corrective hand movement was defined using a variant of the extrapolation method applied to the velocity of the hand in the *x* direction (i.e., the direction of the perturbation) (Oostwoud Wijdenes et al. 2014). We first corrected the *x* velocity in individual change trials. For each block and target change time, we aligned all trials to the time that the target change occurred (in change trials) or the time that the target change would have occurred (in no-change trials). We averaged the *x* velocity in no-change trials to the central target across trials to obtain a ‘baseline’ velocity trace over time, and subtracted this baseline from the *x* velocity in each individual trial. Next, for each trial we obtained the two points at which the additional *x* velocity reached 30% and 70% of the first peak in velocity, and fitted a line through these points and all data points in between. The onset of the reach correction was defined as the time where this line crossed zero. In some trials, the increase in *x* velocity was not approximately linear in the interval between the 30% and 70% peak velocity points, or there was no clear, single peak in *x* velocity, resulting in a poor fit. We therefore discarded trials in which the *R*^2^ of the fit was below 0.95 or in which the extrapolated onset occurred before the target change (2% of trials). Note that trials in which the initial target appeared on the left or the right (and did not change location) were not used in the analysis.

#### Hypotheses and statistical analysis

We designed a set of double-step tasks to determine the effect of target and cue characteristics on the temporal coordination of corrections of the eye and hand. We hypothesized that, with respect to the classic double-step task, corrective saccade latency would increase in the placeholder task, because of the necessity to identify which of the items is the displaced target (Oostwoud Wijdenes et al. 2014; Smeets et al. 2016), as well as in the overlap task, as latency increases are a well-known effect of stimulus overlap (Saslow 1967). We predicted a greater increase in corrective saccade latency in the line and arrow cue tasks, potentially even until after the cursor hit the target, due to the lack of visual stimuli triggering an immediate saccade to the new target. We also hypothesized that corrective saccade latencies would be shorter when online cursor feedback was provided compared to when endpoint feedback was provided, to allow the earliest possible use of visual feedback to help guide the incoming cursor (de Brouwer et al., 2018). If the reach and saccadic system show strong temporal coupling–or shared computation and initiation of corrective responses–we further expect a constant delay between reach and saccade correction latencies across tasks, with high correlations within individuals. Weak coupling–or independent computation and/or initiation of corrective responses–would result in low correlations within individuals, and potentially in varying delays between corrective responses across tasks.

For each of the latencies, we computed the median value per subject, task and feedback condition for no change trials to the central target (20 repetitions), and per subject, task, feedback condition, target change time, and target change direction for change trials (10 repetitions per trial type). These median values were averaged across left and right target change directions, resulting in four values for each combination of task and feedback. To rule out any delays in the corrective saccade due to a saccade refractory period when the target changes during or immediately after the saccade, we assessed the effect of change time on the latency of corrections in the classic double-step task. On average, the target change was visible on the screen 86 ms after saccade onset, 54 ms after reach onset, and 57 ms after the hand passed the midway point, or 242, 343, and 506 ms after target appearance, respectively. Although there was a significant effect of change time on corrective saccade latency, with the shortest latencies when the change occurred after reach onset, there was no significant difference between latencies when the change occurred after saccade onset and when the change occurred midway during the reach (i.e., with a long delay after the initial saccade). This suggests that factors other than the saccade refractory period determined the corrective saccade latency. We therefore averaged the data across change times, calculating the mean and within-subject standard error of the mean (Cousineau 2005) for each combination of task and feedback.

Statistical analyses were performed in R using the rstatix package (v 0.7.0). We performed 5 (task) × 2 (feedback) repeated measures ANOVAs, applying a Greenhouse-Geisser correction when the assumption of sphericity was violated, and we calculated generalized eta squared effect sizes 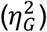. Significant main effects were followed up by pairwise comparisons with Bonferroni correction for multiple comparisons.

## Results

The main goal of this study was to determine the influence of the characteristics of the visual target or cue that indicates a sudden change in movement goal, on the timing of rapid corrective responses of the eye and hand. An additional goal was to determine the effect of feedback of the hand on the timing of these corrections. In the following sections, we will first show task and feedback effects on correction latencies of the eye and hand. We then report how the temporal relation between eye and hand corrections was affected by task and feedback.

### Initial reaction times

Regardless of task and feedback condition, each double-step trial started with a saccade toward the initial (central) target that was followed by an initial reach toward the central target. To provide insight into the timeline of events in our task, we report the mean reaction times of these initial movements, calculated per task and feedback condition, in Table S1. Briefly, the mean reaction times ranged between 143 and 162 ms for initial saccades, and between 197 and 260 ms for initial reach movements. Importantly, all correction latencies in our main analyses were assessed relative to the target change, which was triggered by the execution of the saccade or reach. Therefore, any effects of task and feedback on corrective latencies are independent of any effects on initial reaction times, and we did not test the effects of task or feedback on these initial reaction times. The short initial reaction times suggest that participants were anticipating the appearance of the target, complicating the interpretation of these values. The Pearson correlation coefficients between saccade and reach reaction times, calculated separately for each participant and then averaged across participants, ranged between 0.39 and 0.59 (Table S1), indicating moderate temporal coupling during the initial movement phase.

### Effects of task and feedback on movement corrections

#### Latencies of corrective saccades and reach corrections

In response to the change in movement goal, both the eye and the hand rapidly initiated a correction in all conditions. Examples of these corrections and their timing are shown in Figure 2 for a representative participant, in blocks with online cursor feedback. Figure 2A shows that the initial straight-ahead reach trajectory was followed by a correction to the left or right, with later target changes (brighter colors) eliciting later trajectory adjustments requiring a greater change in reach direction. The eye and hand started correcting at around the same time (Fig. 2B), and the corrective saccade was of much shorter duration than the reach correction.

**Figure 2.**
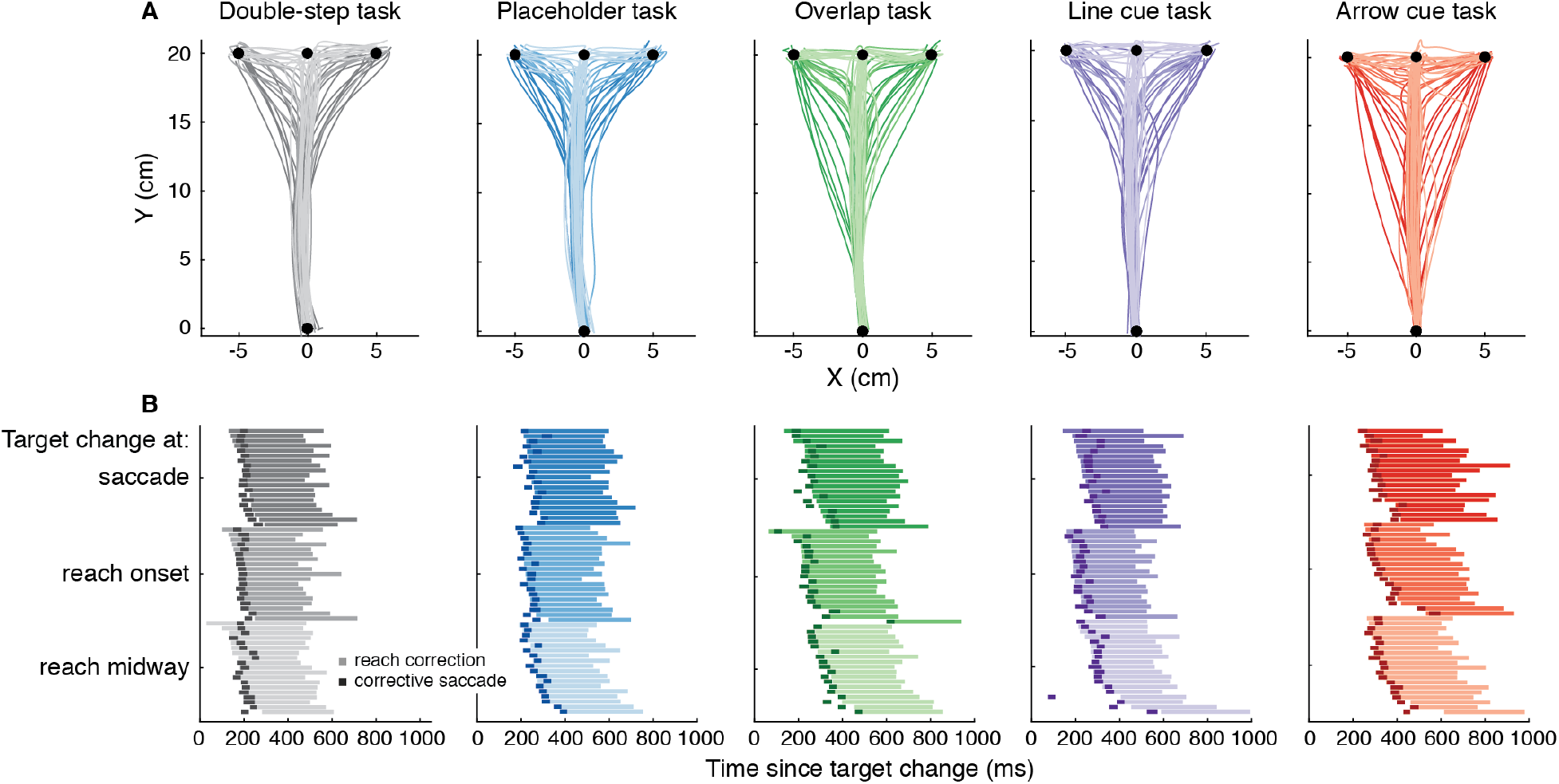
Eye and hand corrections in response to a change in movement goal in individual trials of an example participant, in tasks with online cursor feedback. A) Hand trajectories. Each trajectory is a single trial, with the brightness indicating the target change time as indicated in the legend in B. Note that the results were averaged across target change times (see Methods). B) Timing of corrections relative to the target change. Reach corrections are shown by the longer horizontal bars (one for each trial), with the brightness indicating the target change time. Corrective saccades are shown by the short dark-colored bars. For each target change time, trials are sorted according to the latency of the reach correction.

Averaging across participants revealed a clear task-dependency of correction latencies (Fig. 3A, B). Figure 3A shows that corrective saccade latencies (relative to the time of target change) were modulated by the characteristics of the visual target or cue that indicated the target change. Latencies were shortest in the classic double-step task (*M*=191 ms), longer in the placeholder task (*M*=244 ms), longer again in the overlap and line cue tasks (*M*=271 and 271 ms), and longest in the arrow cue task (*M*=305 ms). This observation was reflected in a significant main effect of task in a 5 (task) × 2 (feedback) ANOVA (*F*(4,60)=67.5, *p*<0.001, 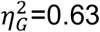). Post-hoc pairwise comparisons confirmed significant differences between all tasks (*p*<0.01), except between the overlap task and the line cue task (*p*=1.000; all *p*-values for pairwise comparisons here and below were adjusted with a Bonferroni correction). Corrective saccade latencies did not differ between blocks with online cursor feedback and blocks with endpoint cursor feedback (main effect *F*(1,15)=0.2, *p*=0.647, 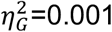). There was no significant interaction between task and feedback (*F*(4,60)=1.5, *p*=0.220, 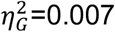).

**Figure 3.**
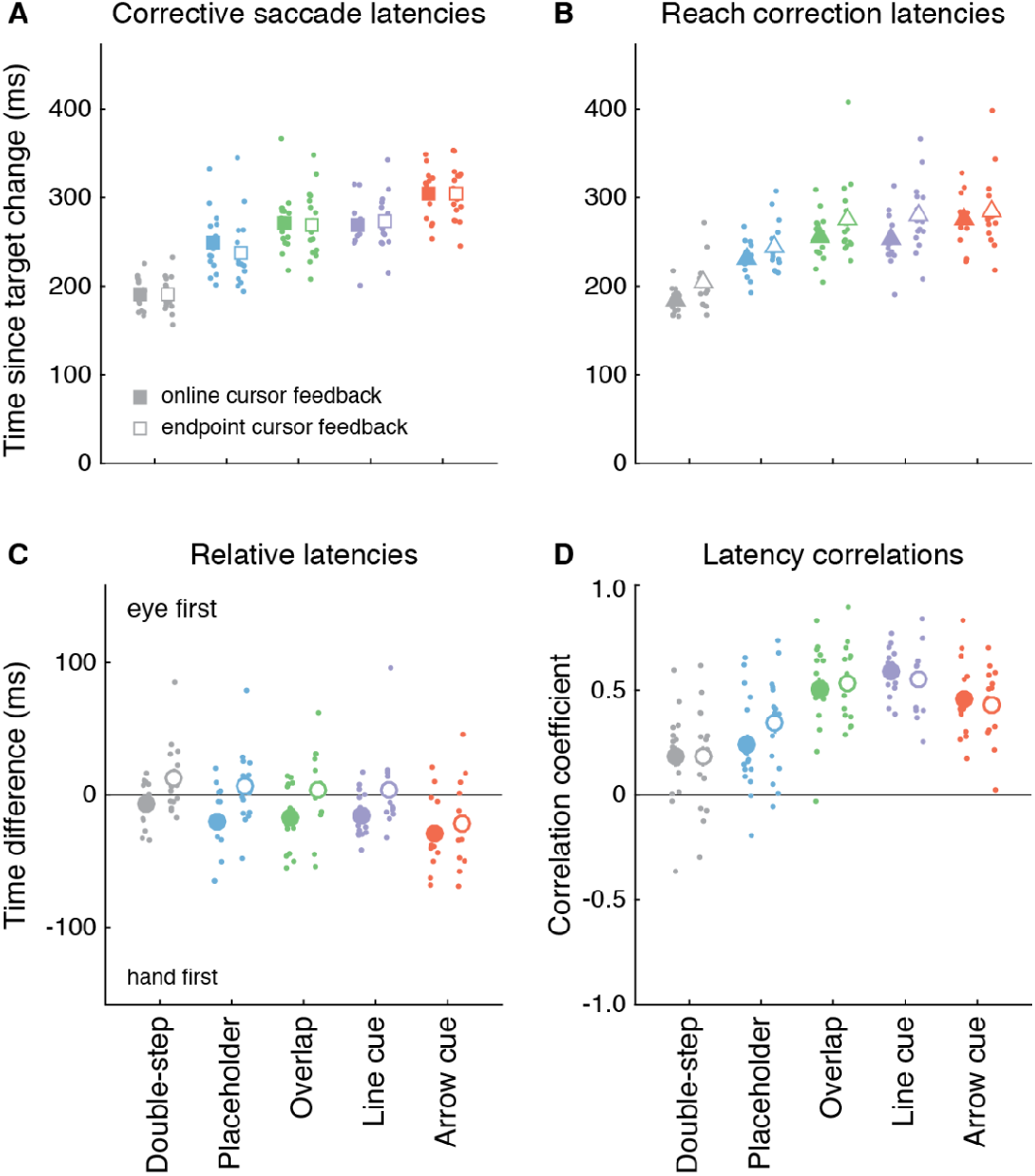
Timing of eye and hand corrections in response to a change in movement goal, as a function of task and cursor feedback. A) Latencies of corrective saccades relative to the target change. Each small dot represents a participant and each square represents the average across participants (*n*=16). B) As A, but showing reach correction latencies. C) Differences between corrective saccade latencies and reach correction latencies, with positive values indicating that the eye started correcting first, and negative values indicating that the hand started correcting first. D) Correlations between corrective saccade and reach correction latencies. For all panels, colors represent different tasks, filled symbols represent online cursor feedback and open symbols represent endpoint feedback only. Error bars represent ± one within-subjects standard error of the mean [note that these are often hidden behind the symbol showing the average value].

Reach correction latencies were also dependent on the characteristics of the visual target or cue that indicated the target change (Fig. 3B). Reach correction latencies were shortest in the double-step task (*M*=194 ms), longer in the placeholder task (*M*=238 ms), and longest in the overlap (*M*=265 ms), line cue (*M*=266 ms) and arrow cue (*M*=280 ms) tasks. This was reflected in a significant main effect of task (*F*(4,60)=57.7, *p*<0.001, 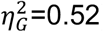), with significant pairwise differences between all tasks (all *p*<0.001), except between the overlap, line cue, and arrow cue tasks (*p*>0.05). In addition, reach correction latencies were shorter with online cursor feedback (*M*=240 ms) than with endpoint cursor feedback (*M*=258 ms; main effect of feedback *F*(1,15)=9.2, *p*=0.008, 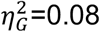). The effect of feedback was significant for all tasks (*p*<0.05) except the arrow cue task (*p*=0.375), as revealed by pairwise comparisons following up on the significant interaction between task and feedback (*F*(4,60)=3.2, *p*=0.018, 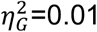).

In summary, corrective saccade latencies were longer in the modified double-step tasks than in the classic double-step task, in line with our expectations. Reach correction latencies followed a similar pattern to saccade latencies, with the exception that there was no further increase in latency for the arrow cue task. We also found that while corrective saccade latencies were not influenced by the presence of online cursor feedback, reach corrections started earlier when online cursor feedback was provided. Knowing that both corrective saccade latencies and reach correction latencies were affected by the task, the next question is how the relative timing and the correlation between saccade and reach corrections were influenced by the task.

#### Temporal coordination of movement corrections

To determine the influence of task on the temporal coordination of corrections of the eye and hand, we calculated the relative latencies of corrective saccades and reach trajectory corrections, as well as the correlations between latencies. Figure 3C shows that the hand often started correcting before the eye. The relative latencies of corrections of the hand and eye showed a significant effect of task (*F*(4,60)=9.8, *p*<0.001, 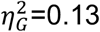) and feedback (*F*(1,15)=15.3, *p*=0.001, 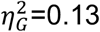), as well as a significant interaction (*F*(4,60)=3.4, *p*=0.014, 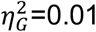). First, the lead of the hand was greater in the arrow cue task (*M*=25 ms) than in the other four tasks (all *p*<0.001). The relative latencies in the double-step, placeholder, overlap, and line cue tasks (*M*=-3, *M*=7, *M*=6 and *M*=6 ms, respectively) did not differ significantly from each other (all *p*>0.05). Second, the lead of the hand was greater when the hand cursor was visible during the reach (*M*=18 ms) than when the cursor was only shown at the end of the reach (*M*=−1 ms). Pairwise comparisons to unpack the interaction showed that this difference was significant for all tasks (all *p*<0.01) except the arrow cue task (*p*=0.080).

As an additional measure of the temporal coordination of corrections, we calculated Pearson correlation coefficients between corrective saccade and reach correction latencies (Fig. 3D). These correlations showed a significant effect of task (*F*(4,60)=18.0, *p*<0.001, 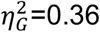) but not feedback (*F*(1,15)=0.3, *p*=0.623, 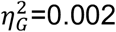), and no significant interaction (*F*(4,60)=1.1, *p*=0.343, 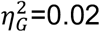). Correlations were lower in the double-step task (*M*=0.18) than in the overlap (*M*=0.52, *p*<0.001), line cue (*M*=0.57, *p*<0.001), and arrow cue tasks (*M*=0.44, *p*<0.001). Correlations were also lower in the placeholder task (*M*=0.29) than in the overlap (*p*=0.003), line cue (p<0.001), and arrow cue tasks (*p*=0.026), and correlations were higher in the line cue task than in the arrow cue task (*p*=0.020).

In summary, the hand generally started correcting its trajectory in response to a change in movement goal before a corrective saccade occurred. Although in some cases the relative delays between corrections of the hand and eye were around zero, the longer neuromuscular delays and higher inertia of the arm (Gribble et al. 2002) imply that even in these cases, the neural signal to execute a correction occurred earlier for the hand than for the eye (see Discussion). Taken together with the results on absolute latencies above, the greater hand-eye delay in the arrow cue task compared to the other tasks was the result of the corrective saccade being delayed. Further, the greater lead of the hand when online cursor feedback was provided resulted from an earlier response of the hand. Finally, correlations between correction latencies differed between tasks, with lower correlations in the classic double-step task and placeholder task than in the overlap, line cue, and arrow cue tasks.

## Discussion

This study aimed at determining the effects of visual target and cue characteristics on temporal eye-hand coordination when hand movements had to rapidly correct for a change in movement goal location, while the eyes were free to move. To date, few studies have investigated online corrections of goal-directed reach movements while also measuring eye movements. In addition, the task demands in these studies were limited to simple target step paradigms. We found that across stimulus and task conditions, participants initiated a corrective saccade in addition to correcting the hand trajectory. We report the following key findings: First, the correction latency in eye and hand depended on visual and cognitive processing demands of the task. For example, task versions that contained a cue indicating the change in movement goal triggered longer-latency corrections as compared to task versions in which participants simply followed the target when it stepped to a new location. Second, eye and hand corrections were not initiated simultaneously: the hand started correcting before the eye. The hand-eye correction delay was greatest when the target change was signaled by a symbolic cue. Third, correction latencies of the eye and hand were weakly correlated, within individuals and tasks, in target step tasks, and moderately correlated in tasks where the target change was indicated by an additional target or cue, requiring an intentional correction. Fourth, online visual feedback of the hand cursor, compared to endpoint feedback only, sped up the initiation of online corrections of the hand trajectory but did not affect the latency of corrective saccades. Together, our findings show that the timing and coordination of rapid movement corrections of the eye and hand depend on the visual and cognitive demands of the change in movement goal. These findings provide important insights into the role of stimulus and task demands during rapid movement corrections, and will be discussed in detail below.

Our first key finding is that the latencies of both corrective saccades and reach corrections depend on the processing demands of the task. Previous studies already showed that visual target characteristics can affect reach correction latencies in response to a target change (Kozak et al. 2019; Veerman et al. 2008). Here, we show that corrective saccade latencies are also affected by the visual as well as cognitive demands of the task. Specifically, consistent with our hypothesis, corrective saccade latencies increased, relative to those in the classic double-step task, in a placeholder task where all three possible targets were shown and the target swapped position with one of the placeholders, and further increased in an overlap task where the new target was shown in addition to the initial target. Corrective saccade latencies also increased relative to the classic double-step task when the target change was indicated by a spatial (line) cue, and further increased when the target was indicated by a symbolic (arrow) cue. The latencies of corrections of the reach trajectory showed a similar pattern, but without a further increase in latency in the arrow cue task. Note that in all of our tasks, reach correction latencies were slower than commonly reported. This is most likely a result of the extra mass and inertia of the robotic manipulandum, but could also have been caused by relatively low contrast of stimuli on the screen (Veerman et al. 2008). Nevertheless, the task-dependency of latencies emphasizes the importance of visual characteristics of the target and the requirements of the task for the timing of movement corrections.

Our second key finding is that corrections in eye and hand are not initiated simultaneously, and–as the absolute latency–the relative latency of corrections depends on the visual and cognitive processing demands of the task. Whereas many previous studies reported that the eye typically leads the hand when initiating a movement (Bowman et al. 2009; Land and Hayhoe 2001; Neggers and Bekkering 2000; Prablanc et al. 1979), here we show that the opposite can be true when performing a correction, as Abekawa and colleagues (2014) have previously shown for corrective movements in response to a target displacement (see also Gritsenko et al. 2009). Although the relative latencies between corrections of the hand and the eye were around zero in most of the tasks in the current study, it is important to note that even in cases where corrections of the eye and hand were detected at around the same time from their velocities, the longer neuromuscular delay and greater inertia of the arm and hand imply that the neural signal to execute a correction was sent earlier to the arm muscles than to the eye muscles. Specifically, the neuromuscular delay of the hand, obtained by measuring surface electromyographic (EMG) responses at the first dorsal interosseous (index finger) muscle following transcranial magnetic stimulation of the motor cortex, is as short as 20 to 30 ms (Werhahn et al. 1994), and the electromechanical delay (the delay between EMG activity and movement) ranges anywhere between 40 and 120 ms (Challis 2021), resulting in a total neuromechanical delay of 60 to 150 ms (see also Del Vecchio et al. 2018). For saccades, a neuromechanical delay of 25 ms following electrical stimulation of the monkey frontal eye fields has been reported (Robinson and Fuchs 1969), and this value is likely comparable in humans. Thus, corrective saccade onset has to occur at least 35 ms ahead of corrective hand movement in order for the saccade neural signal to precede the neural signal for the hand. This was only the case for one out of 16 participants, who showed relative latencies between 45 and 100 ms (eye first) when only endpoint cursor feedback was provided. Across all participants, the average lead of the hand ranged between −3 and 25 ms, suggesting that the neural signal for the hand preceded that for the eye by at least 32 to 60 ms, on average.

The finding that the reach correction is initiated before the corrective saccade indicates that the initial correction of the reach must be based on an approximate computation of the visual movement goal location (Franklin et al. 2016), since high-resolution retinal information as well extraretinal information of the new target position does not become available until the eye lands on the new target. By contrast, the later part of the correction could be more refined, as shown for trajectory corrections in response to cursor displacements (Cross et al. 2019). Further, the delay of the corrective saccade was greater in the arrow cue task compared to the other tasks, as a result of a further increase in corrective saccade but not reach correction latencies. In this particular task, the eye appeared to linger, likely reflecting the time it took to process the shape information of the visual cue, requiring the involvement of the ventral visual stream (Goodale and Milner 1992). Moreover, in the arrow cue task, the cue was presented centrally, akin to tasks involving endogenous visual spatial attention. Such tasks typically involve slower and more sustained shifts of attention. By contrast, the line cue task involved an exogenous cue, which is known to trigger faster, transient shifts of visual spatial attention (Busse et al. 2008; Carrasco 2011). The time course of events in our tasks reflects the differences between endogenous and exogenous visual spatial attention. However, it is somewhat surprising that the extra processing time in the symbolic cue task affected the eye, but did not cause a further increase in reach correction latencies. This might suggest that corrective responses of the hand can be initiated before cognitive processing is complete.

As another aspect of temporal eye-hand coordination, we showed that, within individuals and tasks, correction latencies of the eye and hand are weakly to moderately correlated. Our third key finding is that these correlations were higher for tasks that involved a slower, intentional or voluntary correction (overlap and cue tasks) than in tasks that involved a faster, more automatic correction (classic double-step and placeholder tasks). This is similar to the findings reported by Sailer and colleagues (2000) for direct goal-directed actions. We propose that the higher correlations in more demanding tasks might result from an overlap in the latency distribution of eye and hand movements for slower, voluntary saccades but not for faster saccades, or might result from the presence of a ‘bottleneck’ in information processing that affects both the oculomotor and the limb-motor system (though this would not explain the greater hand-eye delay in the arrow cue task). Some studies have interpreted temporal correlations in favor of a common command for eye and hand movements (e.g., Herman et al. 1981). A series of recent studies on eye-hand corrections used a version of a double-step task in which participants were instructed to cancel their response to the initial target and direct their eyes and hand to the new target instead. Using a drift-diffusion framework, the results were better explained by a common accumulator for the eye and hand than two separate, interacting accumulators (Gopal and Murthy 2015, 2016). When participants performed a dual task in which an instructed eye-only or hand-only response had to be substituted by a movement with both the eyes and hand when a tone was presented with the target, responses were most often–but not always–best predicted by the separate, interacting accumulator model (Jana et al. 2017). Based on the finding that the common and separate command model could explain subsets of trials with different behavior in each task, the authors concluded that participants can use both models, with the frequency determined by task context (Jana et al. 2017; Jana and Murthy 2018). It is important to note however, that in these tasks the eyes and hand always made the same movement from the start location to the target, whereas in our tasks the hand corrected in-flight and the eyes performed an additional, corrective saccade. Furthermore, in the current study temporal correlations between movement corrections were weak to moderate and differed across tasks. This suggests that the initiation of eye and hand corrections is driven by separate movement commands served by different neural circuitry (de Brouwer et al. 2021).

Our fourth key finding is that online visual feedback on the hand (cursor) location, as compared to endpoint feedback only, sped up the initiation of hand movement corrections (Reichenbach et al. 2009), resulting in a greater lead of the hand. Corrective saccade latencies did not differ between blocks with online and endpoint cursor feedback This was in contrast to our hypothesis that online cursor feedback would (also) reduce the latency of corrective saccades, given the fact that directing the eyes to the reach target improves monitoring of the reach trajectory in peripheral vision (de Brouwer et al. 2018), providing an important reason to move the eyes to the new target location as soon as possible. Presumably, cursor feedback increases certainty about the location of the hand (Acerbi et al. 2017; Izawa and Shadmehr 2008), and therefore accelerates the correction, but this facilitation does not transfer to the eye.

With respect to the neural mechanisms underlying movement corrections, previous studies on online corrections of reaching movements have emphasized the role of the posterior parietal cortex (Desmurget et al. 1999; Pisella et al. 2000), as well as the possibility of a subcortical pathway guiding both the eye and the limb when performing corrections (Day and Brown 2001; Reynolds and Day 2012) (see also Cross et al. 2019; Pruszynski et al. 2010). The exact contributions of these areas are still unknown (for reviews see Archambault et al. 2015; Gaveau et al. 2014). Reynolds and Day proposed that a subcortical circuit, involving the superior colliculus, allows for very fast, automatic responses, while a cortical pathway drives slightly slower but more flexible responses. Although our results cannot provide conclusive evidence about the pathways involved in online corrections, it is highly likely that both these pathways contribute to the corrective responses in our task. We propose that the subcortical circuit plays a greater role in the classic double-step task, where responses were fastest and most automatic. The other tasks required more visual and cognitive processing, therefore responses in these tasks were most likely mediated by a cortical circuit, resulting in slower responses. Further, the involvement of the ventral visual stream can explain the additional increase in latencies in the arrow cue task. Our results also suggest that while (initial) visual and cognitive processing of the new target or cue might be shared, corrective responses for the eye and hand are initiated by different neural circuits. It is possible that, given the instruction to reach to the target as fast as possible, the response of the hand is prioritized over the response of the eye, and saccades might be actively suppressed until the response of the hand is initiated (Mrotek and Soechting 2007).

In summary, we designed a set of double-step tasks and showed that the correction of the reach trajectory in response to a target change is accompanied by a corrective saccade. Both the absolute and relative latencies of the corrective saccade and reach correction depended on the visual characteristics of the target change. Corrections were initiated later when the task required more visual and cognitive processing, and the hand typically started correcting before the eye, especially when the change was indicated by a symbolic cue. Our results highlight that the hand and eye are coordinated in a flexible manner that is suited for the task at hand (see also Fooken et al. 2021). Our findings also provide a framework for discussing latency differences obtained across tasks and studies and emphasize the importance of taking stimulus and task conditions into account when assessing eye-hand coordination. This is especially important when aiming to generalize laboratory results to the more visually and cognitively demanding conditions of real-world tasks.

## Supplemental Data

Publicly available DOI for Figshare data: https://doi.org/10.6084/m9.figshare.14781738.

## Acknowledgements

We would like to thank Hannah Brown for help with data collection. This work was funded by an NSERC Discovery Grant and Accelerator Supplement (RGPIN 418493) awarded to Miriam Spering.

